# Visualizing somatic alterations in spatial transcriptomics data of skin cancer

**DOI:** 10.1101/2022.12.05.519162

**Authors:** Limin Chen, Darwin Chang, Bishal Tandukar, Delahny Deivendran, Raymond Cho, Jeffrey Cheng, Boris C. Bastian, Andrew L. Ji, A. Hunter Shain

**Affiliations:** Department of Dermatology, University of California San Francisco; Department of Immunology, H. Lee Moffitt Cancer Center; Department of Pathology, University of California, San Francisco; Helen Diller Family Comprehensive Cancer Center, University of California, San Francisco; Department of Dermatology, Department of Oncological Sciences, Black Family Stem Cell Institute, Tisch Cancer Institute, Icahn School of Medicine at Mount Sinai

## Abstract

Tools to visualize genetic alterations within tissues remain underdeveloped despite the growth of spatial transcriptomic technologies, which measure gene expression in different regions of tissues. Since genetic alterations can be detected in RNA-sequencing data, we explored the feasibility of observing somatic alterations in spatial transcriptomics data. Extracting genetic information from spatial transcriptomic data would illuminate the spatial distribution of clones and allow for correlations with regional changes in gene expression to support genotype-phenotype studies. Recent work demonstrates that copy number alterations can be inferred from spatial transcriptomics data^1^. Here, we describe new software to further enhance the inference of copy number from spatial transcriptomics data. Moreover, we demonstrate that single nucleotide variants are also detectable in spatial transcriptomic data. We applied these approaches to map the location of point mutations, copy number alterations, and allelic imbalances in spatial transcriptomic data of two cutaneous squamous cell carcinomas. We show that both tumors are dominated by a single clone of cells, suggesting that their regional variations in gene expression^2^ are likely driven by non-genetic factors. Furthermore, we observe mutant cells in histologically normal tissue surrounding one tumor, which were not discernible upon histopathologic evaluation. Finally, we detected mono-allelic expression of immunoglobulin heavy chains in B-cells, revealing clonal populations of plasma cells surrounding one tumor. In summary, we put forward solutions to add the genetic dimension to spatial transcriptomic datasets, augmenting the potential of this new technology.

## Introduction

The human body is a mosaic of genetically distinct cells^3^ – the result of somatic alterations steadily accumulating in each cell throughout life. Most mutations are neutral and do not affect cellular phenotypes. However, some mutations reduce cellular fitness, contributing to the process of aging^4^, while other mutations increase their fitness, which can ultimately lead to cancer^5^.

Resolving the spatial distribution of mutant cells in diseased and normal tissues will shed light on the earliest phases of tumor evolution. Somatic mutations can mark clonal populations of partially transformed cells that maintain normal histopathological phenotypes (e.g. “field” cells). Moreover, tumors of later stages often are composed of genetically distinct subclones. Defining the boundaries of these subclones can help determine the relative contributions of genetic and non-genetic factors that influence heterogeneity in gene expression. The spatial distribution of somatic alterations within tissues is typically mapped using *in situ* hybridization, *in situ* sequencing, or in some instances immunohistochemistry^6–10^. However, these assays are limited in their scope and the types of somatic alterations that can be detected. As an alternative approach, we investigated whether somatic alterations could be visualized in spatial transcriptomic data.

RNA-sequencing is mainly used to quantify transcript levels but can detect single nucleotide variants in expressed transcripts. Tools to reveal single nucleotide variants in spatial transcriptomic data have not been developed.

Copy number changes at the DNA level can also be inferred from RNA-sequencing data. A program known as inferCNV was developed to derive copy number alterations from single-cell RNA-sequencing data^11^, and it was recently applied to spatial transcriptomic datasets^1^. Another program, known as STARCH^12^, also can infer copy number information from spatial transcriptomic data. Both software packages calculate moving averages of gene expression across the transcriptome to produce copy number estimates. However, as we detail below, this strategy requires further optimization because it is prone to errors when adjacent genes are coordinately regulated.

Taken together, approaches to map somatic alterations in tissues remain underdeveloped. To address this need, we put forward solutions to visualize point mutations, copy number alterations, and allelic imbalance in spatial transcriptomic datasets.

## Results

### Selection of cases

We used publicly available exome sequencing data with matched spatial transcriptomics data of two cutaneous squamous cell carcinomas to detect and visualize the spatial distribution of somatic alterations^2^. Skin cancers have a high mutation burden and thus are well suited for this endeavor. The spatial transcriptomics data had been generated with the 10X Visium platform. Throughout the manuscript, we refer to the patients, from which these tumors were derived, as patient 4 and patient 6 (their original names in Ji *et al*^2^).

The tumors from patient 4 and patient 6 had 121-fold and 214-fold DNA-sequencing coverage over the exome with a computationally inferred 12.1% and 20.7% neoplastic cell content, respectively. This coverage and tumor cell content are sufficient to detect point mutations that are fully clonal as well as larger subclones.

Coverage in spatial transcriptomics data is typically measured by the number of unique molecular identifiers (UMIs) per spot. The tumor from patient 4 had ~15,000 UMIs per spot, and the tumor from patient 6 had ~1,300 UMIs per spot. While the coverage for tumor 6 was low, it allowed us to assess the lower limit of coverage at which somatic alterations can be visualized.

### Defining a reference set of somatic alterations from DNA-sequencing data

We analyzed the exome sequencing data to identify reference sets of somatic alterations for each carcinoma^13^. The mutation burdens were high −24.5 mut/Mbase and 12.2 mut/Mbase for patients 4 and 6, respectively (Table S1) – with strong UV signatures, as is typical for cutaneous squamous cell carcinoma. We found several mutations in known driver genes. The tumor from patient 4 had a *TP53^E285K^* mutation, and the tumor from patient 6 had *NOTCH1^E2071K^, MTOR^S2215F^, TP53^P278L^, CHUK^V587M^,* and *CDKN2A^R7*/D23A^* mutations. The tumor from patient 4 had no discernible copy number alterations or allelic imbalances, whereas the tumor from patient 6 had several arm-level gains and losses with allelic imbalance patterns that were generally concordant with the underlying copy number alterations (Fig. S1). Taken together, the high burden of point mutations, primarily attributable to UV radiation, and spectrum of driver mutations were consistent with previous genetic characterization of cutaneous squamous cell carcinoma^13^.

### Visualization of point mutations in 10X Genomics Visium data

The Visium platform captures and sequences transcripts from the poly-A tail, limiting mutation detection to those near the 3’ end of expressed genes (see an example in Fig. S2). Given these constraints, we detected 36 mutations (4.5% of all mutations from the exome DNA-sequencing data) in the spatial transcriptomics data from the patient 4 tumor and 17 mutations (3.7% of the total mutation count) from the patient 6 tumor (Table S1).

Next, we mapped sequencing reads that spanned these mutation sites across the tissue of each tumor (Fig. 1). We considered a spot with one or more mutant reads to harbor tumor cells, and we considered a spot unlikely to contain tumor cells if it had 5 or more reference reads without any reads of the mutant alleles. The higher threshold to call tumor-free spots reflects the possibility that the wild-type allele can be sampled from heterozygous mutations (Fig. 1). A dermatopathologist previously annotated the tumor regions within each biopsy, blinded to the results of our genetic analyses. The regions histopathologically annotated as tumor were mostly covered by spots containing mutant reads (Fig. 1).

**Figure 1.**
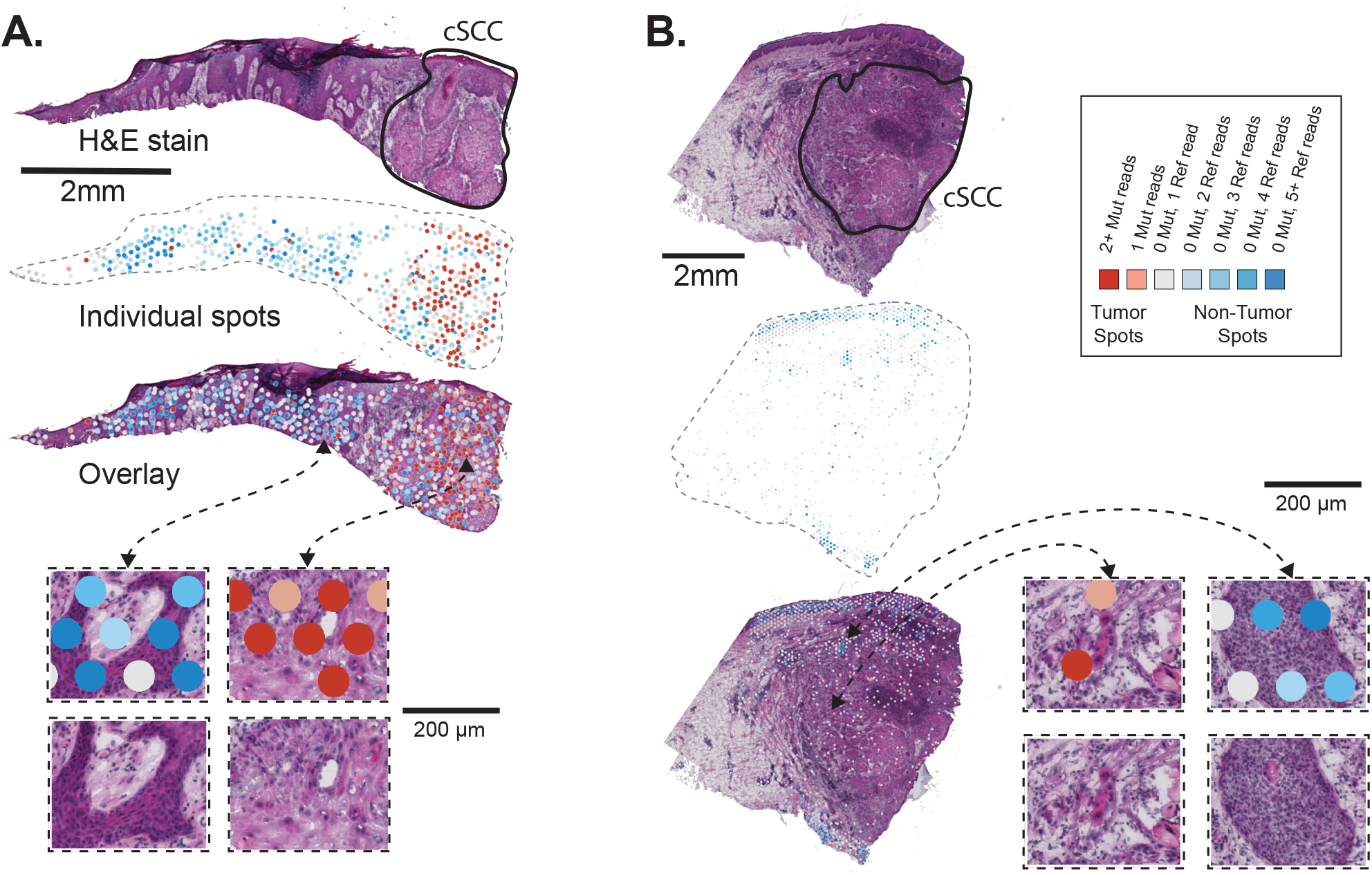
Somatic point mutations are detectable in spatial transcriptomics data. **A-B.** H&E stains are shown for cutaneous squamous cell carcinomas from patients 4 and 6 of Ji *et. al*. Cell (2020). The sections underwent histopathologic assessment, and areas of cutaneous squamous cell carcinoma (cSCC) are circled as shown. DNA sequencing was performed on these tumors to call somatic mutations. Spots from the spatial transcriptomics arrays are colored based on the presence or absence of sequencing reads mapping to the mutant or reference alleles over somatic mutation sites. Note the enrichment of mutant spots in tumor areas.

Unexpectedly, we observed spots with mutant reads in histologically normal tissue from patient 4. These spots were situated in an area of actinic keratosis, a precursor of squamous cell carcinoma as well as in a region of reactive epidermal hyperplasia interposed between the actinic keratosis and squamous cell carcinoma (Fig. 2A).

**Figure 2.**
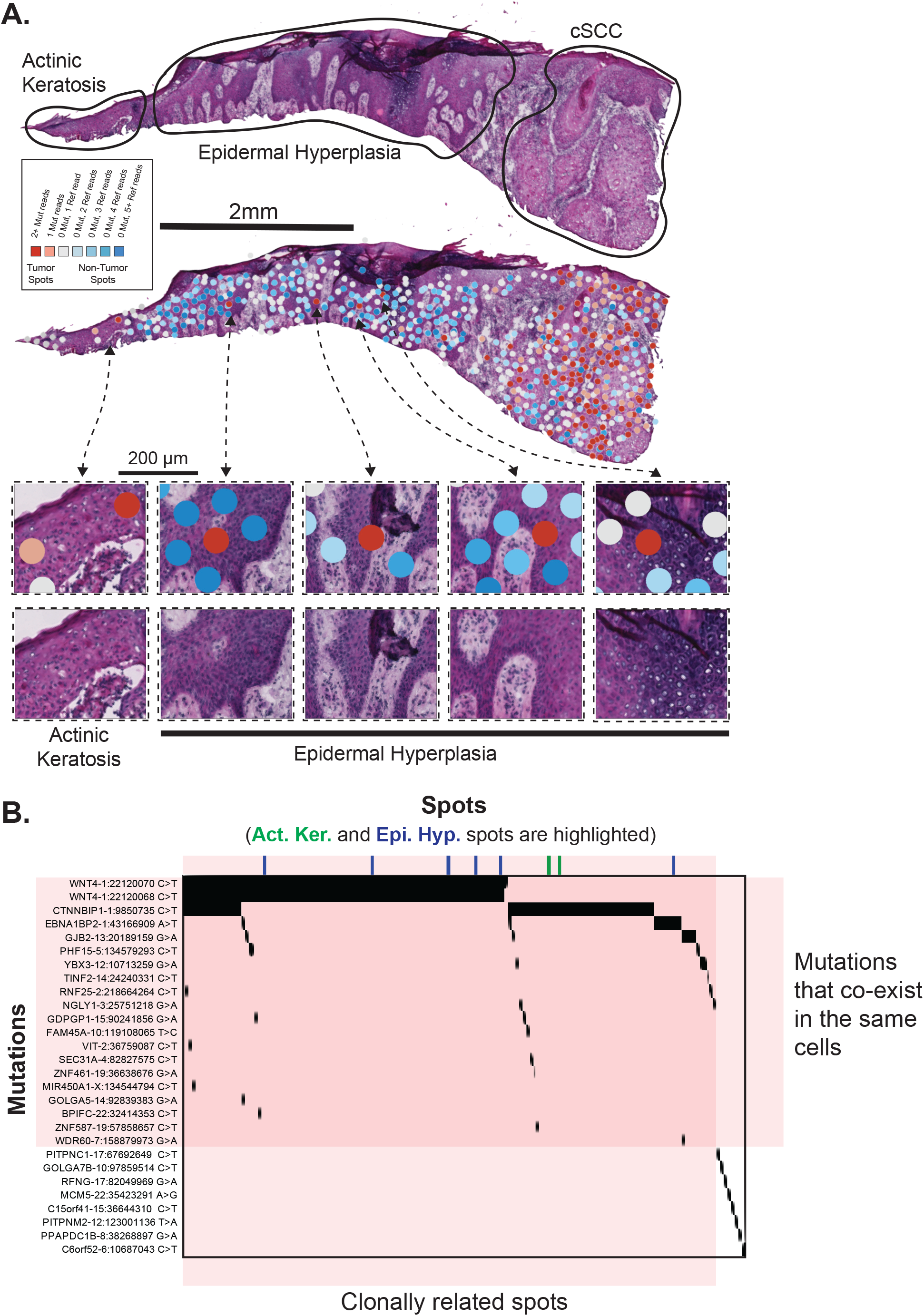
Somatic mutations are observed in histologically normal cells outside of the cutaneous squamous cell carcinoma. **A.** H&E stain of the tumor from patient 4 with mutant spots annotated as described in Figure 1. Zoomed insets focus on spots with somatic mutations that are outside of the cSCC region. **B.** A tiling plot shows the distribution of mutations (rows) across spots (columns) from the spatial transcriptomic data. Black tiles register when a mutation was present in a given spot. White tiles had too little coverage to make a definitive call. There were no tiles for which a mutation could conclusively be called as absent (0 mutant reads and 5+ reference reads). The tiling plot only displays the subset of spots and mutations for which mutant reads could be detected in the spatial transcriptomics data.

We considered the possibility that mutant reads detected in the histologically benign tissue might have originated from RNA molecules in the neighboring tumor tissue that diffused to other spots during hybridization. To assess the possibility of such leakage, we inspected the total read counts in the spots that were not covered by any tissue (Fig. S3). While spots covered by tissue had a median of 16,709 reads, spots outside of the tissue only had a median of 213 reads. The presence of sequencing reads in spots not covered by tissue indicates that some level of diffusion of mRNA does occur, but mRNA abundance is two orders of magnitude higher over tissue spots. Next, we inspected the mutant read counts in areas outside of the tissues and found trace reads with mutations (0.15 mutant reads per mm^2^). The density of mutant reads in the non-cancerous tissue areas (1.45 mutant reads per mm^2^) was ~10-fold higher than the density of mutant reads outside of the tissue areas altogether (Fig. S3B). Taken together, diffusion of mRNA was unlikely to account for the number of mutant reads in the actinic keratosis or the area of reactive epidermal hyperplasia.

The clonal relationship between all spots with mutations is shown in Fig. 2B. The black tiles indicate a specific mutation (row) that was present in a given spot (column). The empty tiles indicate that the mutation was not detected, however this does not imply that the mutation was truly absent. Definitively determining the absence of mutations was limited by the low coverage over the mutant bases. 98.8% of the empty tiles had no reads and the remaining 1.2% had fewer than 5 reads.

The pattern of mutations across the tissues implies that most of the mutant spots are clonally related, regardless of their spatial localization. Many mutations co-existed in at least a subset of spots (Fig. 2). The co-occurrence of mutations in the same spot indicates that they co-exist within the same cells. Normal skin cells can have a high burden of somatic mutations^14,15^ of their own, but the mutations observed in the normal skin of patient 4 were shared with the neighboring squamous cell carcinoma, suggesting a common ancestry between their cells (Fig. 2).

We did not observe evidence of subclones in the tumor. A small number of somatic mutations did not overlap with other mutations and could not be assigned to a clone (Figure 2B, see bottom right of tiling plot). These mutations may mark spots from unrelated clones of cells, but they affected genes with low transcript levels, suggesting that the mutations were likely missed in the other spots. Deeper DNA-sequencing and/or deeper spatial transcriptomic-sequencing would eventually reveal subclones of cells, but our analysis suggests there is a dominant clone of mutant cells in this tumor biopsy.

### Visualization of copy number alterations in 10X Genomics Visium Data

While levels of gene expression are affected by many variables, it is possible to infer the DNA copy number of the underlying genes from RNA-sequencing data by averaging transcript levels of multiple adjacent genes in a sliding window along the chromosome^11,16,17^. This strategy reduces the variability in expression of individual genes to instead reveal the changes in gene expression, across a larger segment of the genome, that typically accompany copy number alterations. Our laboratory expanded upon this approach with CNVkit-RNA^18^, which gives transcripts more weight for copy number calling when their gene expression shows a high correlation with copy number changes of the underlying genes in The Cancer Genome Atlas project.

We used CNVkit-RNA to infer copy number information from individual spots, and it detected arm-level copy number alterations over a subset of spots that paralleled those seen in the exome sequencing data (Fig. 3A). To establish cutoffs for calling copy number alterations for a given spot, we calculated a score to reflect how similar the copy number profile of each spot was to the copy number profile inferred from the bulk-cell DNA-sequencing of the tumor (see methods). To determine whether these scores were statistically significant, we calculated the same score on permuted data to establish a null distribution of scores. The observed scores were, on average, higher than the scores from the permuted data, indicating that there was an enrichment of spots with true copy number signals matching the tumor’s DNA copy number profile (Fig. S4). Spots with copy number alterations tended to overlie regions histopathologically annotated as tumor (Fig. 3B).

**Figure 3.**
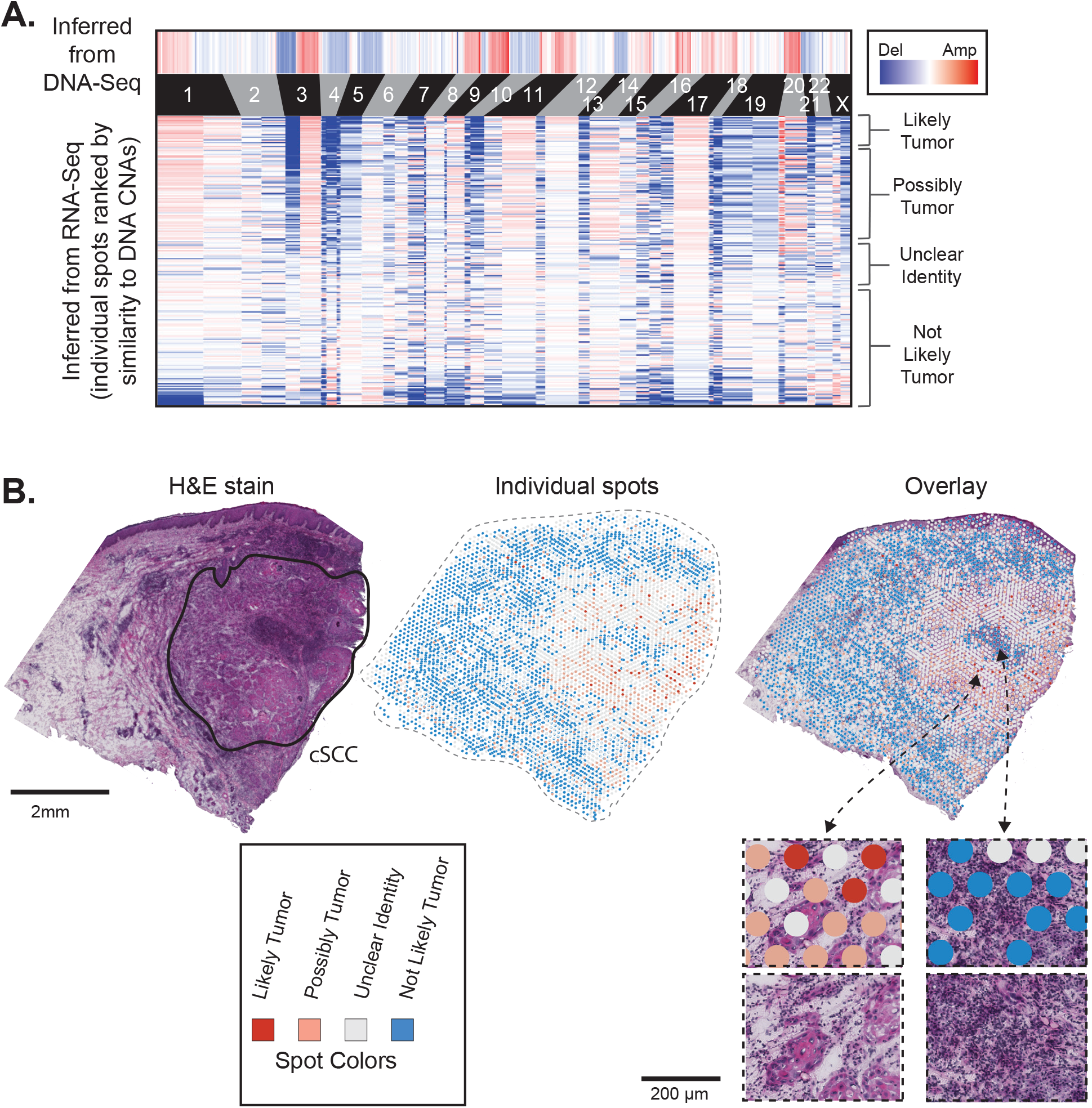
Copy number alterations are detectable in spatial transcriptomics data. **A.** Copy number alterations (CNAs) were inferred from DNA-Sequencing data (top heatmap) and from RNA-sequencing data of individual spots (lower heatmap). Spots (rows in the lower heatmap) are ranked ordered by the similarity of their copy number profiles to the DNA copy number alterations and classified into groups, ranging from “Likely Tumor” to “Not Likely Tumor” (See Figure S3 for more information on groupings). **B.** An H&E stain of the patient 6 cutaneous squamous cell carcinoma is shown with the main tumor region circled. Individual spots are colored as indicated in panel A. Spots are shown by themselves and overlaying the the H&E image.

We also benchmarked CNVkit-RNA^18^ against InferCNV^11^. After running InferCNV, we identified a similar set of copy number aberrations as a recent study^1^, which also used InferCNV on the patient 6 data. The most prominent copy number signals detected by InferCNV were absent from the bulk DNA-sequencing data from this tumor (Fig. S5). InferCNV predicted copy number alterations in genomic regions with clusters of lineage-specific genes. For instance, copy number gains in keratinocytes were predicted over genomic regions containing a cluster of keratin genes, and copy number gains in lymphocytes were predicted over families of immune-related genes (Fig. S5). The most likely explanation is that the moving average of gene expression spiked over these clusters of highly expressed genes, producing false positive copy number calls in tissue areas enriched with certain cell types. By contrast, the weighting algorithm used by CNVkit-RNA did not flag these loci as affected by copy number changes (Fig. 3), in agreement with the patient-matched DNA-sequencing data.

While copy number signals were detectable in spatial transcriptomics data, they were less specific for identifying tumor cells than point mutations. Detection of copy number signals is more susceptible to false positives due to systematic variation in gene expression, as our permutation model shows (Fig. S4). Indeed, some spots predicted to have copy number changes were located outside of the marked tumor area in patient 6 (note the occasional red or pink spots in the stroma surrounding the tumor in Fig. 3B); however, the density of these spots did not exceed the density expected by chance, based on the false discovery rates associated with those spots. The spurious nature of these spots is supported by the absence of spots with point mutations in these areas (Fig. 1). Overall, we believe that point mutations can be used to localize rare populations of mutant cells in spatial transcriptomics datasets (as shown in Fig. 2), whereas copy number alterations are best used to visualize general areas enriched with cells harboring copy number alterations.

### Visualization of allelic imbalance in 10X Genomics Visium Data

Next, we tested whether allelic imbalance could be detected in spatial transcriptomic data. Heterozygous SNPs were identified from the bulk-cell DNA-sequencing data of each patient’s normal tissue. We also counted the number of reads mapping to each allele in the tumor’s DNA-sequencing data and designated the more abundant allele as the “major” allele. Most of the tumor genome showed marginal differences in allele counts from the tumor’s DNA-sequencing data, resulting in arbitrary assignments, but there were some contiguous genomic regions with clear-cut imbalances (e.g. chromosome 3q of patient 6, Fig S1B).

We plotted the ratio of reads mapping to the major:minor allele for each SNP and from each spot (Fig. 4A-B). If a SNP shows mono-allelic expression, then all reads would map to either the major or minor allele, evident in the scatterplot as having a 1:0 or 0:1 ratio of reads. Mono-allelic expression was most common in poorly expressed genes, as would be expected due to the higher variability when sampling low numbers of reads. There was a notable exception (Fig. 4B), discussed below, in which a highly expressed SNP showed monoallelic expression.

**Figure 4.**
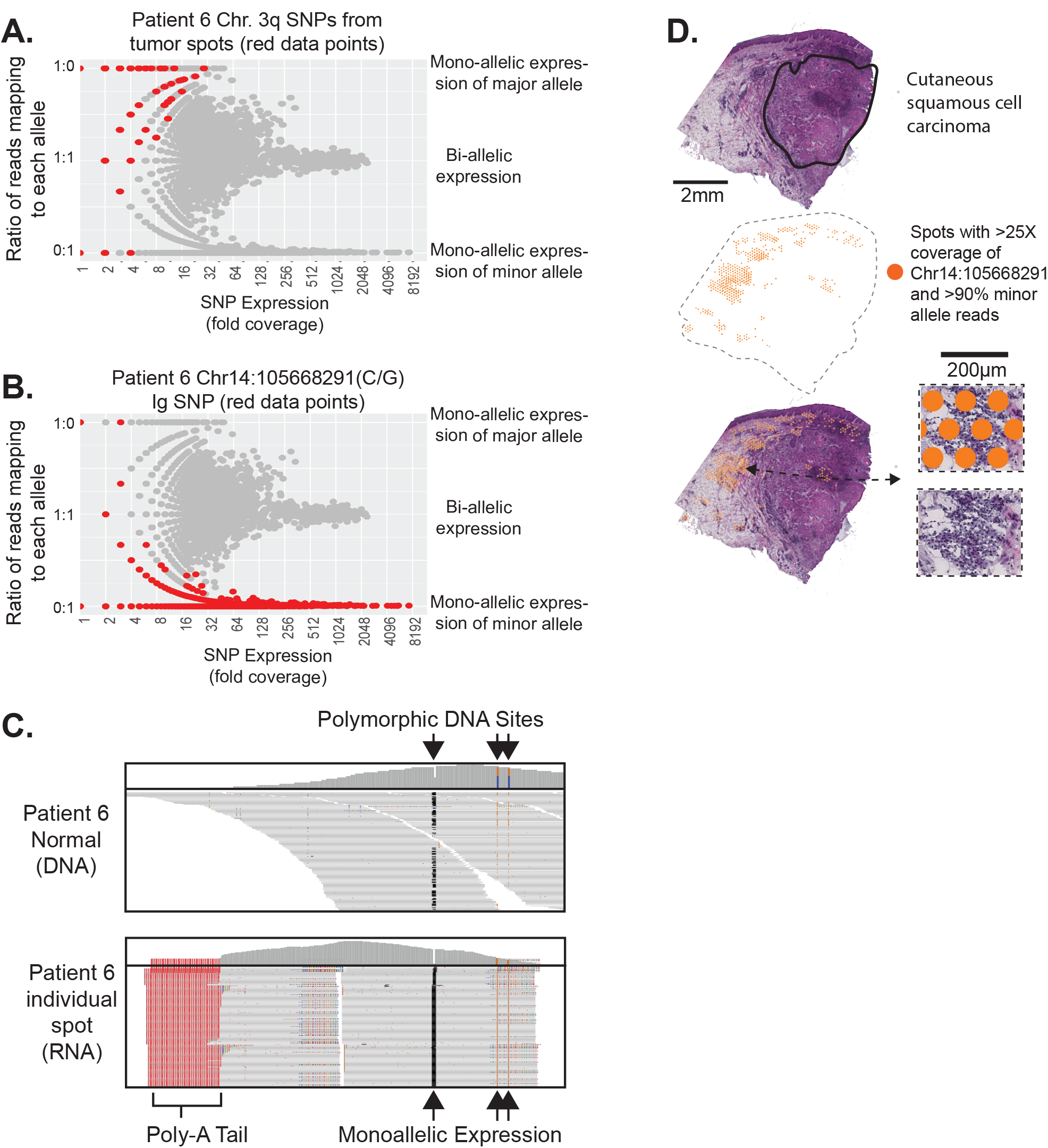
Allelic imbalance is detectable in spatial transcriptomics data. **A-B.** We counted the number of reads mapping to each allele of heterozygous SNPs from each spot. To identify SNPs with mono-allelic expression, we plotted the fraction of reads mapping to each allele as a function of total read coverage. As expected, the reads from genes with low expression often came from one allele, however, genes with high expression tended to express both alleles with a notable exception highlighted in panels B-D. Panel A highlights SNPs from chromosomal arm 3q of tumor spots, for which there was loss of heterozygosity evident in tumor DNA-sequencing data (see Figure S1B). Panel B highlights a SNP from the immunoglobulin heavy chain (IGH) locus at chromosome 14q32 in which expression favors one allele. **C.** DNA- and RNA-sequencing read alignments surrounding the IGH SNP. Note heterozygosity in the DNA-sequencing data, while the RNA-sequencing data shows monoallelic expression. **D.** Spots with greater than 25X coverage and >90% of reads mapping to the minor allele of the IGH SNP are projected onto the tissue, where they tend to overlie immune cells.

As a benchmark, we measured allelic expression of heterozygous SNPs on the X-chromosome of patient 4, who was female. We observed mono-allelic expression of X-chromosome SNPs (Fig. S6A), consistent with the expected silencing patterns that result from inactivation of one X-chromosome. X-chromosome inactivation randomly occurs during development, resulting in mosaic silencing patterns in tissues^19^. While it is possible that a spot could overlie 2 cell populations in which different X-chromosomes were inactivated, previous studies showed that the typical clone size of cells with shared X-chromosome inactivation is much larger than the spot size of a Visium array, mainly due to the early stage of development in which X-chromosome inactivation occurs^20^. Concordantly, neighboring spots also tended to express the same allele, supporting the notion that mosaic clones occupy significant volumes in adult tissues (Fig. S6B). X-chromosomal SNPs in the *XG* and *RPS4X* genes were outliers in that they retained bi-allelic expression (Fig. S6C-D), but this was expected as these genes are known to escape X-chromosome inactivation^19,21,22^.

We next measured allelic imbalance in the spatial transcriptomics data of heterozygous SNPs on chromosome 3q in tumor spots from patient 6. The DNA-sequencing data detected allelic imbalance in this region, likely caused by an underlying copy number gain. Consistent with this observation, the corresponding major alleles were preferentially expressed in the spots overlying tumor cells for these SNPs (Fig. 4A). Allelic imbalance in other chromosomal regions was too subtle to be reliably detected for both tumors.

Finally, we explored allelic imbalance in an unbiased manner. There were several heterozygous SNPs mapping to the immunoglobulin heavy chain locus, which were highly expressed, exclusively from one allele (Fig. 4B, C). Immunoglobulin genes undergo somatic rearrangement during maturation of B cells, and after re-arrangement, the un-rearranged allele is silenced (an observation that has been termed “allelic exclusion”^23–25^). Allelic exclusion ensures that the mature B-cell produces a single antibody. The spots with high levels of mono-allelic expression, localized to the periphery of the tumor, in regions with an increased density of immune cells (Fig. 4D). The full-length sequence of the mRNA from the immunoglobulin heavy and light chain would be needed to assemble the VDJ rearrangement and delineate the precise clonal relationship between the different areas of B-cells. However, the allelic exclusion, observed here, suggests that a clonal population of B-cells encircled the tumor.

## Conclusions

Our work builds on pioneering studies demonstrating that somatic copy number alterations can be visualized in spatial transcriptomics data^1,12^. Here, we establish that three types of genetic alterations – somatic mutations, somatic copy number alterations, and germline polymorphisms – are detectable in spatial transcriptomics data. Among these, somatic point mutations provide the highest specificity in marking cells with underlying alterations. However, a major limitation to the detection of point mutations is the need for sufficient coverage over the mutant base pair. Detection of copy number changes is possible at higher sensitivity but at comparatively lower resolution.

Germline SNPs provide additional information. Recognizing imbalance between the alleles requires a sufficient number of reads covering the SNP. SNPs in highly expressed genes, such as immunoglobulin genes, satisfy this requirement. For other SNPs, broad regions with loss-of-heterozygosity can be revealed by integrating coverage over polymorphisms in cis along the same chromosome. As an example, we were able to distinguish haplotypes over chromosomal arm 3q in one sample, due to the imbalance in DNA-sequencing reads from the patient’s tumor. In doing so, we confirmed that the major allele is predominately expressed in tumor cells.

Taken together, these three genetic readouts provide complementary types of information to enrich the analysis of spatial transcriptomics data. We used this information to reveal tumor cells extending into histologically normal skin adjacent to a cutaneous squamous cell carcinoma. It is not uncommon for cutaneous squamous cell carcinomas to recur after surgical removal^26^. Our data suggests that recurrent tumors could arise from a reservoir of cells that persist in normal-appearing tissue. We also used allelic imbalance data to identify what is most likely to be a clonal population of B-cells surrounding one tumor. Knowing the clonal structure of immune cells, in addition to their gene expression profiles, provides valuable information to inform our understanding of interactions between tumor cells and the adaptive immune system^27^.

Finally, our data suggests that each tumor was dominated by a single clone. The mutant allele frequencies of the somatic mutations had a unimodal distribution; the copy number alterations were of similar amplitude; and we did not observe genetic differences between the cells in different areas of these tumors. Ji *et. al*. previously revealed gene expression heterogeneity in these tumors – particularly when comparing cells at the leading edge versus the interior of the tumors^2^. It is most likely that these spatially defined gene expression programs are driven by non-genetic factors. Our observations do not preclude the possibility that genetic heterogeneity exists at higher resolution, such as in individual cells, but it is unlikely that the cells at the leading edge of cutaneous squamous cell carcinoma, which constitute ~1/3^rd^ of these tumors, are genetically distinct from the interior of the tumor, despite their unique gene expression patterns.

In summary, current workflows to analyze spatial transcriptomic data have not fully unlocked the potential of this technology. We show that genetic alterations can be detected and visualized in spatial transcriptomics data, and we anticipate these workflows will facilitate genotype-phenotype studies by adding the genetic dimension to spatial transcriptomic datasets.

## Acknowledgements

We wish to acknowledge research support from: American Cancer Society Research Scholar Grant, Tracy and Guy Jacquier cSCC Research Fund, Mt. Zion Health Fund, UCSF Resource Allocation Program, University of California Cancer Research Coordinating Committee, LEO Foundation, United States Department of Defense, the UCSF Department of Dermatology, NIH K08CA263187, and NIH R01CA265786. We would also like to acknowledge computational research support from the Computational Biology and Informatics Shared Resource through the Helen Diller Family Comprehensive Cancer Center at UCSF.

## Methods

### Code and data availability

All scripts are available on GitHub: https://github.com/limin321/stmut. We include Youtube tutorials walking through the main analyses performed by these scripts: https://www.youtube.com/playlist?list=PLK-4mLUJI-Xr1NJiMq2887D8BpMveBuX2. Raw data is available on GEO: accession number GSE144237 and GSE144239. Source data for each figure is available at the Github page above.

### Assembling Exome and Spatial Transcriptomic sequencing data

Whole exome DNA-sequencing data and spatial transcriptomics data was generated by Ji and colleagues and made publicly available as previously described^2^. Briefly, after isolation of genomic DNA, it was prepared for sequencing, and libraries were enriched with exome baits (Agilent SureSelect Human All Exon V6). Separate tumor sections were placed on 10X Visium arrays (slide serial number: V19T26-101), hybridized, and prepared for sequencing according to manufacturer’s protocols. There were two replicates (sequential sections of tissue) from each tumor biopsy, which were processed for spatial transcriptomics. The data from each replicate was processed in parallel and integrated as described below. The DNA-sequencing data is available at NCBI Gene Expression Omnibus (GEO) (accession number GSE144237). The spatial transcriptomics data is available from NCBI GEO database (accession number GSE144239).

### Calling somatic alterations from DNA-sequencing data (Related to TableS1 and Figure S1)

We previously performed a meta-analysis of exome sequencing studies covering cutaneous squamous cell carcinoma where we called somatic point mutations, copy number alterations, and allelic imbalances from these two tumors, among others^13^. A candidate list of somatic point mutations was generated with MuTect (v4.1.2.0) by comparing the tumor sequencing alignments to patient-matched reference alignments. This list was filtered to generate a final list of somatic mutations, as described (https://github.com/darwinchangz/ShainMutectFilter). The point mutation calls are available as part of this manuscript in Table S1. Copy number was inferred with CNVkit^28^, and a candidate list of germline polymorphisms was generated with FreeBayes by identifying variants when comparing the normal sequencing alignments to the reference genome. A final list of germline, heterozygous SNPs was inferred by identifying those variants that overlapped with known 1000 genome SNPs and which had variant allele frequencies between 40-60%. The raw copy number calls (cnr and cns files produced by CNVkit) and a list of germline, heterozygous SNPs (patient4_hg38_SNPs.txt and patient6_hg38_SNPs.txt files produced with our filtering) are available in the GitHub repository associated with this manuscript (https://github.com/limin321/stmut/tree/master/VisualizingSomaticAlterations/DNAseqResourceFiles/).

### Aligning spatial transcriptomics sequencing data to the transcriptome

Fastq files were aligned to the hg38 genome using the Space Ranger pipeline (spaceranger-1.3.0) by 10X genomics, as previously described^2^. This pipeline produces a single bam file with sequencing reads aggregated from all spots. Next, we split this bam file into individual bam files for each spot using the subset-bam script by 10X Genomics (https://github.com/10XGenomics/subset-bam). This script outputs hundreds to thousands of individual bam files, depending on the number of spots, each with sequencing reads matching the barcode tag for individual spots.

### Visualizing somatic point mutation reads in spatial transcriptomics data (Related to Figures 1, 2, and S2)

At this point, somatic point mutations had been identified from DNA-sequencing data, and the sequencing alignments from the spatial transcriptomics data had been split into individual bam files based on the spatial barcode tag in each read, resulting in hundreds of bam files per spatial transcriptomics run (one bam file per tissue-covered spot). We next used the mpileup function from samtools to count mutant and reference reads over the somatic mutation sites (defined from the DNA-sequencing data) in each of the bam files corresponding to an individual spot. Our script loops through each somatic mutation site from each bam file, and is available on GitHub (https://github.com/limin321/stmut) along with an instructional video walking through them on YouTube (https://www.youtube.com/watch?v=pvs_b1ALygA). After counting individual mutant sites from each spot’s bam file, we summarized the mutant allele and reference allele counts within each spot.

Spots were combined into the following groups, as indicated in the legend of Figure 1: spots with 2 or more mutant reads, spots with 1 mutant read, and spots with no mutant reads. Spots with only 1 mutant read were considered likely to be tumor spots because the probability of a false positive is equivalent to the error rate during the sequencing process, which is low. Nevertheless, these spots were manually inspected to eliminate obvious artifacts. We removed a total of three spots (all from patient 6 replicate 2) that had issues. These mutant reads were in the incorrect orientation and/or had numerous mismatches throughout the read length. Including them would not have affected the conclusions of this manuscript.

Spots with 0 mutant reads were further subdivided, as indicated in Figure 1, based on the number of reference reads, ranging from 1 reference read to 5 or more reference reads. Since most somatic point mutations are heterozygous, tumor cells can produce reference reads when the wild-type allele is sampled during sequencing. Therefore, a small number of reference reads does not indicate that the spot in question had no tumor cells; however, the probability that there are no tumor cells increases as the number of reference reads increases in the absence of mutant reads.

Once spots were grouped, as described above, we imported their barcodes into the Loupe browser (10X Genomics) and selected customized color schemes to visualize the spots from each group, as shown in the legend of Figure 1. Two images were exported — a “spots only” image and an “H&E only” image. The tumors from patient 4 and 6 had two replicates each. To merge the data from the replicates, we subtracted the background from the “spots only” image and overlayed the spots from both replicates onto the “H&E only” image of each tissue in Adobe Illustrator.

### Quantifying background signals on a Visium Array (Related to Figure S3)

As part of the Space Ranger workflow, there is a step in which the user defines the spots overlying tissue. Removing non-tissue spots improves gene expression clustering and principal component analyses by eliminating datapoints without true signal; however, we sought to use the read coverage over non-tissue spots as a proxy of background signals that may arise from diffusion of mRNA during hybridization.

Towards this goal, we ran the Space Ranger workflow a second time and selected all spots as overlying tissue. UMI counts per gene per spot were exported using the mat2csv command (a function within the Space Ranger software distribution), producing a table from which we could count the number of reads per spot. A heatmap showing the number of reads per spot is shown in Figure S3A (note the exponential scale). We also split the aggregate bam file into individual bam files using the 10X Genomics subset-bam script and counted the number of somatic mutant reads per spot, as described above. A Loupe projection showing the localization of mutant spots is shown in Figure S3A.

We grouped spots into three categories – non-tissue spots, benign tissue spots, and tumor tissue spots as shown in Figure S3. After grouping, we calculated the total number of reads per spot, the number of mutant reads per spot, and the surface area of spots from each group. A table summarizing these statistics is shown in Figure S3B. We specifically highlight the number of mutant reads per square millimeter in benign tissue versus non-tissue areas in the bar graph to the right of Figure S3B. The error bars correspond to 95% confidence intervals (Poisson-test).

### Inferring somatic copy number alterations in spatial transcriptomics data (related to Figure 3)

Copy number was inferred from each spot of the patient 6 tumor biopsy. We did not attempt copy number analyses of the patient 4 tumor because the DNA-sequencing data did not predict there to be any alterations.

To infer copy number alterations from each spot, we first generated a matrix of unique molecular identifier (UMI) counts from each gene/spot using the mat2csv command from the spaceranger software distribution. We combined the data from replicates 1 and 2 of patient 6 into a single matrix to be processed together.

We used the import-RNA command^18^ in the CNVkit package^28^ to convert the UMI counts to logarithmic ratios of gene expression (centered based on the median signal within the dataset itself). This command also filtered out genes with poor expression across the spots, and it assigned a weight to each gene, upweighting genes that are better able to provide copy number information. The weight is an important feature of CNVkit that differentiates it from other methods to infer copy number from RNA-sequencing data. Briefly, CNVkit calculates a weight for each gene that is proportional to that gene’s correlation between expression and copy number from cancer genome atlas data – the net effect is that genes whose expression is known to concord with copy number in independent datasets are given more weight. CNVkit further modifies the weight based on the variability of gene expression and the absolute level of gene expression within the dataset being analyzed – genes with relatively stable expression and relatively higher expression are given more weight. Collectively, a gene with a high weight can provide a more reliable copy number estimate than a gene with a low weight.

The standard approach to inferring copy number information from RNA-sequencing data is to calculate a moving average of expression over a window of genes^11,16,17^. We borrowed this concept, but we also sought to incorporate the weights, assigned by CNVkit. When we originally developed the import RNA command for CNVkit, we used pre-existing segmentation algorithms that were able to incorporate the weight values for each gene^18^. These segmentation algorithms worked well on bulk RNA-sequencing data^18^, however, they did not test well on spatial transcriptomics data because they were originally designed for DNA-sequencing data. Therefore, for this manuscript, we wrote an R-script to calculate the weighted median of expression from genes on the same chromosomal arm (https://github.com/limin321/stmut) along with an instructional video walking through them on YouTube (https://www.youtube.com/watch?v=QIDp9TLICuU), offering arm-level copy number inferences across the genome for each spot.

Before proceeding further, we filtered out spots with no UMIs on 2 or more chromosomal arms. We attempted to rescue these spots by combining data from groups of adjacent spots that had been filtered out. After combining data from adjacent spots that had been filtered out, we re-analyzed the data in a second pass. The groups of combined spots had more reads than the individual spots within each group and therefore were less likely to be filtered out on the second pass. When creating groups of spots, we only combined data from adjacent, contiguous spots. In addition, we only combined data from spots assigned to the same gene expression cluster to prevent combining spots encompassing dramatically different populations of cells. Individual spots were grouped together until their total UMI count exceeded 5000 UMIs – typically 2-10 spots per group. We include our grouping script in the GitHub software distribution: https://github.com/limin321/stmut/.

Next, we re-centered the copy number estimates. When CNVkit generated logarithmic ratios of gene expression, it used the median expression of a gene across all spots as its reference point. Consequently, without re-centering, a copy number alteration would appear as a low-level gain (or loss) in tumor spots and a concomitant low-level loss (or gain) in non-tumor spots. Non-tumor spots were inferred by their histology and the gene expression clusters for which they were assigned. For instance, spots assigned to “cluster 4”of patient 6 replicate 1 using the 10X Space Ranger software expressed immune-related genes and tended to overlie lymphocytes – thus, they were classified as non-tumor spots. Any spot with an ambiguous identity was left out of the reference pool. Once we settled upon a reference, we calculated the median copy number signal over each chromosomal arm from the reference pool and subtracted this signal from all spots.

### Comparing the copy number alterations inferred from spatial transcriptomics data to the copy number alterations inferred from patient-matched bulk-cell DNA-sequencing data (Related to Figure S4)

After inferring copy number alterations from spatial transcriptomics data, we sought to compare them to the copy number inferences from the matched DNA-sequencing of the tumor. From the DNA-sequencing data of patient 6, we identified gains of 1p, 3q, 8q, 9q, 11q, 14q, 17q, 20p, and 20q as well as losses of 3p, 4q, 5q, 10p, 10q, 13p, 13q, and 21q (Fig. S1B).

We calculated a score to identify the spots with copy number profiles that were more similar to the DNA-sequencing reference point. The score was calculated as the sum of copy number signals over the regions of known gain minus the sum of copy number signals over regions of known deletions. We also weighed the copy number signals so that they were proportional to the number of genes on each arm – this reduced the influence of small chromosomal arms, whose signal often stemmed from a small number of genes and tended to have more variability.

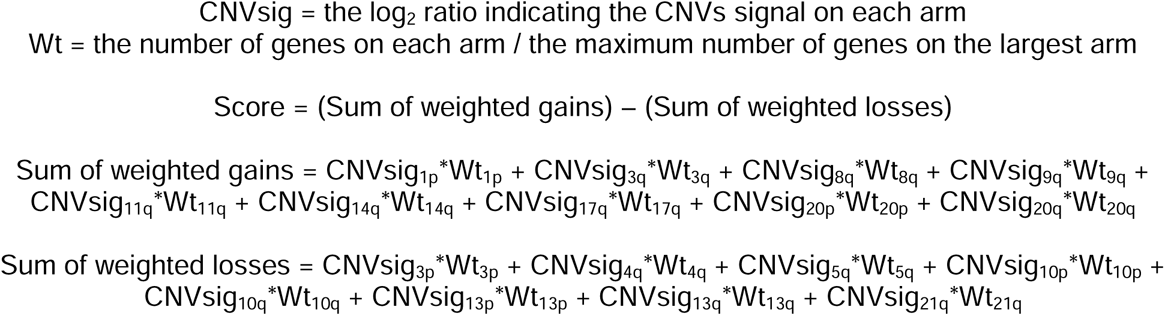

A spot with a copy number profile that is more similar to the DNA-sequencing reference will have a positive score. However, a positive score can arise by random chance. Thus, to better put these scores in context, we permuted the copy number signals from each spot. Permuting the copy number signals from each spot effectively provides a random sampling of copy number alterations that could, in theory, be observed. After permuting the data, we calculated similarity scores on the permuted data to provide a theoretical distribution of scores that could occur by random chance. We produced 138,400 permuted scores (100-fold more datapoints than the observed data, which covered 1384 spots). The histogram of permuted scores and observed scores are shown in Figure S4A. Our permutation script is available in GitHub – https://github.com/limin321/stmut/blob/master/FigTableScripts/FigTables.md#figure-s4.

We further calculated a false discovery rate for each spot. We counted the number of permuted datapoints at a given spot’s score or higher and divided by 100 to normalize for the size of the permuted dataset relative to observed data – this number was considered the number of false positives at a given score. The total positives were counted from the observed data at a given score or higher. The q-value was calculated by dividing the number of false positives by the number of total positives. The spots with FDR values less that 0.5 are shown in Figure S4B.

### Benchmarking copy number inferences against InferCNV (Related to Figure S5)

In addition to generating copy number calls with CNVkit-RNA, we also generated calls using InferCNV^11^. We ran InferCNV under default conditions. A previous study also used InferCNV to make copy number calls on the exact same dataset^1^. In that study, the authors used a reference pool of single-cell RNA-sequencing data from patient-matched normal tissue to center their data. Under the default conditions, our data was centered relative to the median signal within the dataset itself. Given these differences in centering strategies, the amplitude of some copy number alterations differs between our analysis and those from Erickson and colleagues^1^. Nevertheless, the most prominent copy number inferences were similar in both our analysis as well as the Erickson analysis and are shown in Figure S5.

The highest amplitude copy number calls made by InferCNV were not made by CNVkit-RNA, nor were they evident in the copy number inferred from the DNA-sequencing data (Fig. S5). We investigated the genes at the center of each alteration, and we noted that they tended to encode clusters of lineage-specific genes. For example, amplifications were predicted in keratinocyte cell populations over genes involved in keratinization. As another example, amplifications were predicted in immune cells over genes involved in immune functions. Given that these copy number alterations were not observed in the DNA-sequencing data and that they can easily be explained by the high expression of these genes in certain cell types, we suggest that these are most likely false positives.

The main reason why CNVkit-RNA did not make these same calls is because CNVkit-RNA downweighted most lineage specific genes when inferring copy number. Also, CNVkit-RNA only attempted chromosomal arm-level inferences. Of note, the typical spot from this sample only had ~1,300 UMIs, which corresponds to ~700 detected genes (~15 genes per chromosomal arm). Given the sparse gene coverage, we elected to restrict our analyses to chromosomal arm-level inferences.

To be sure, there was a set of copy number alterations inferred in the DNA-sequencing data as well as in the tumor spots by CNVkit-RNA, InferCNV (our analysis), and InferCNV (Erickson *et. al*. analysis). Examples include loss of 3p, gain of 3q, loss of 4q, loss of 5q, gain of 11p, loss of 13, and gain of 20. As such, we believe that InferCNV can be used to detect copy number alterations in spatial transcriptomics data, however, users should be aware of false positives induced by neighborhoods of co-regulated genes.

### Measuring allelic imbalance in spatial transcriptomics data (related to Figure 4 and Figure S6)

To measure allelic imbalance in spatial transcriptomic data, it is imperative to generate a high-quality list of germline heterozygous SNPs to be interrogated. For instance, if a homozygous SNP were mistakenly input into the heterozygous SNP list, then 100% of reads in the spatial transcriptomic data would map to a single allele, implying that mono-allelic expression was occurring. Artifactual SNP calls also pose a challenge and must be removed. The RNA libraries are prepared for sequencing in a different manner than DNA-sequencing libraries, and the RNA reads are aligned to the genome with different software. Consequently, artifactual SNPs, which were called in DNA-sequencing data, will not necessarily be present in RNA-sequencing data, which would, once again, imply mono-allelic expression was occurring. Using a highly specific list of heterozygous SNPs will alleviate these issues, but we nonetheless recommend users to manually inspect sequencing alignments supporting any notable results.

To ensure the quality of our heterozygous SNP calls, we required SNPs to have at least 10-fold coverage in the normal DNA-sequencing data, to have variant allele frequencies between 40-60% mapping to each allele, and to have been observed in the 1000 Genomes Project in more than 1% of participants. The requirement for high coverage in our reference bam as well as the strict range of allowable allele frequencies ensured that the candidate variants from our data were well supported. The requirement that the variant also be observed in greater than 1% of 1K genome participants ensured that the variant had been observed in another high-quality dataset, though we likely missed SNPs that are rare in the general population.

While the heterozygous SNPs were defined from the donor’s normal DNA-sequencing data, we also counted the number of reads mapping to the ref and alt allele in the tumor DNA-sequencing data, and we renamed the more abundant allele in the tumor DNA-sequencing data as the “major allele”. This was a meaningful designation when there was clear-cut allelic imbalance in the DNA-sequencing data. However, for much of the genome, allelic imbalance was not present, or it was too subtle to definitively identify the more abundant allele. Therefore, the “major allele” designation was arbitrary for many SNPs – an assignment based on whichever allele was randomly sampled at greater frequency during DNA sequencing of the tumor.

Once we generated a list of germline heterozygous SNPs, we counted expression of each SNP’s allele in each spot’s bam file using the mpileup command in the samtools software distribution. Our approach to counting reads mapping to each SNP allele was the same as the approach we used to count reads mapping to mutant and wild-type alleles at somatic mutation sites, as described above. The specific scripts related to these analyses are available here: https://github.com/limin321/stmut/blob/master/FigTableScripts/FigTables.md#figure-4 along with an instructional video walking through them on YouTube (https://www.youtube.com/watch?v=diZDaFUahzc).

Most SNPs had no expression mapping to either allele because they did not reside in the sequenced portion of an expressed gene. Nevertheless, SNPs are relatively common, so there were 1772 SNPs from donor 4 and 2071 SNPs from donor 6 with at least one read of coverage over the SNP site in at least one spot. A list of SNPs and their coverage in each spot is available in the GitHub repository here: https://github.com/limin321/stmut/tree/master/ResourceFiles/Figure4SourceData.

For each SNP from each spot, we plotted the fraction of reads mapping to the major allele versus the total coverage. When coverage is low, one would expect a broader spread in allele frequencies, due to random sampling biases and transcriptional bursts^29^, and this is indeed what we observed (Fig. 4). At higher coverage, read ratios tended to stabilize at one-to-one ratios mapping to the major/minor alleles. We used these plots to identify SNPs with disproportionate expression of a single allele. A SNP from the immunoglobulin locus of patient 6 primarily expressed the minor allele (Fig. 4C-E), as discussed in the main text. In addition, two SNPs in *S100A8* of patient 4 primarily expressed the major allele, but we concluded that these were most likely mapping artifacts. We discuss why these were most likely mapping artifacts in the separate section below entitled, “*Mapping artifacts in SNPs from patient 4 (Related to Figure 4)*”.

Coverage over most other SNPs was too low to recognize allelic imbalance in the spatial transcriptomics data. Therefore, we explored allelic imbalance in a hypothesis-driven manner. We identified a region with allelic imbalance over chromosomal arm 3q from the DNA-sequencing data of the patient 6 tumor. No tumor spots from patient 6 had greater than 32X coverage over a heterozygous SNP from this chromosomal arm; however, when we visualized the read distribution of all SNPs from this arm, there was a skew towards reads mapping to the major allele (Fig. 4A).

We also investigated allelic read coverage of SNPs on the X-chromosome of patient 4. Patient 4 was female, and therefore one would expect mono-allelic expression over heterozygous SNPs on the X-chromosome due to inactivation. We observed mono-allelic expression of X-chromosome SNPs (Fig. S6A) for all but two SNPs (Fig. S6C). The two outliers occurred in genes known to escape X-chromosome inactivation, as discussed in the main manuscript.

Of note, the tumor from patient 6 also came from a female donor, and we observed mono-allelic expression for all SNPs on the X-chromosome with coverage. However, read depths were extremely low and coverage across spots was too sparse to perform similar analyses as shown for patient 4 in Fig S6B, D.

### Mapping artifacts in SNPs from patient 4 (Related to figure 4)

In patient 4, there were two SNPs (Chr1:153419253[G/A] and Chr1:153418150[G/A]) that appeared to primarily express the major allele. Upon further inspection, these SNPs are most likely to be mapping artifacts. Both SNPs map to the *S100A8* gene. *S100A8* is one of 24 genes in the S100 gene family, most which cluster on chromosome 1q. The genes in this family are extremely homologous, sharing approximately 50% similarity in amino acid sequences^30^, making it challenging to unequivocally map sequencing reads to the appropriate genes in this family. This challenge is exacerbated by the 3’ sequencing strategy, utilized by 10X Genomics. Sequencing data consists of 120bp single-end reads, but many reads are soft-clipped, reducing their effective length, because they extend into the template switching oligo or the poly-A tail. Considering these challenges, we noted that the reads mapping to the major allele of these SNPs mapped similarly well to other S100 genes. In addition, the Chr1:153419253[G/A] SNP was 112 base pairs away from the poly-A tail, yet there was only 112X coverage over the poly-A tail while there was 67,000X coverage over the SNP site. We did not observe such precipitous drops in coverage over any other gene. Local spikes in read coverage, such as this, are common features of alignment artifacts in RNA-sequencing data. Based on this body of evidence, we determined that further evidence was needed to conclude that monoallelic expression was occurring in the *S100A8* gene.

---

**Table S1. Somatic mutations in tumors from patient 4 and 6.** Point mutations were called as described and annotated with the funcotator tool from genome analysis toolkit (see GATK for description of column headers).

Other figure legends are embedded in the figure files.

**Figure S1.**
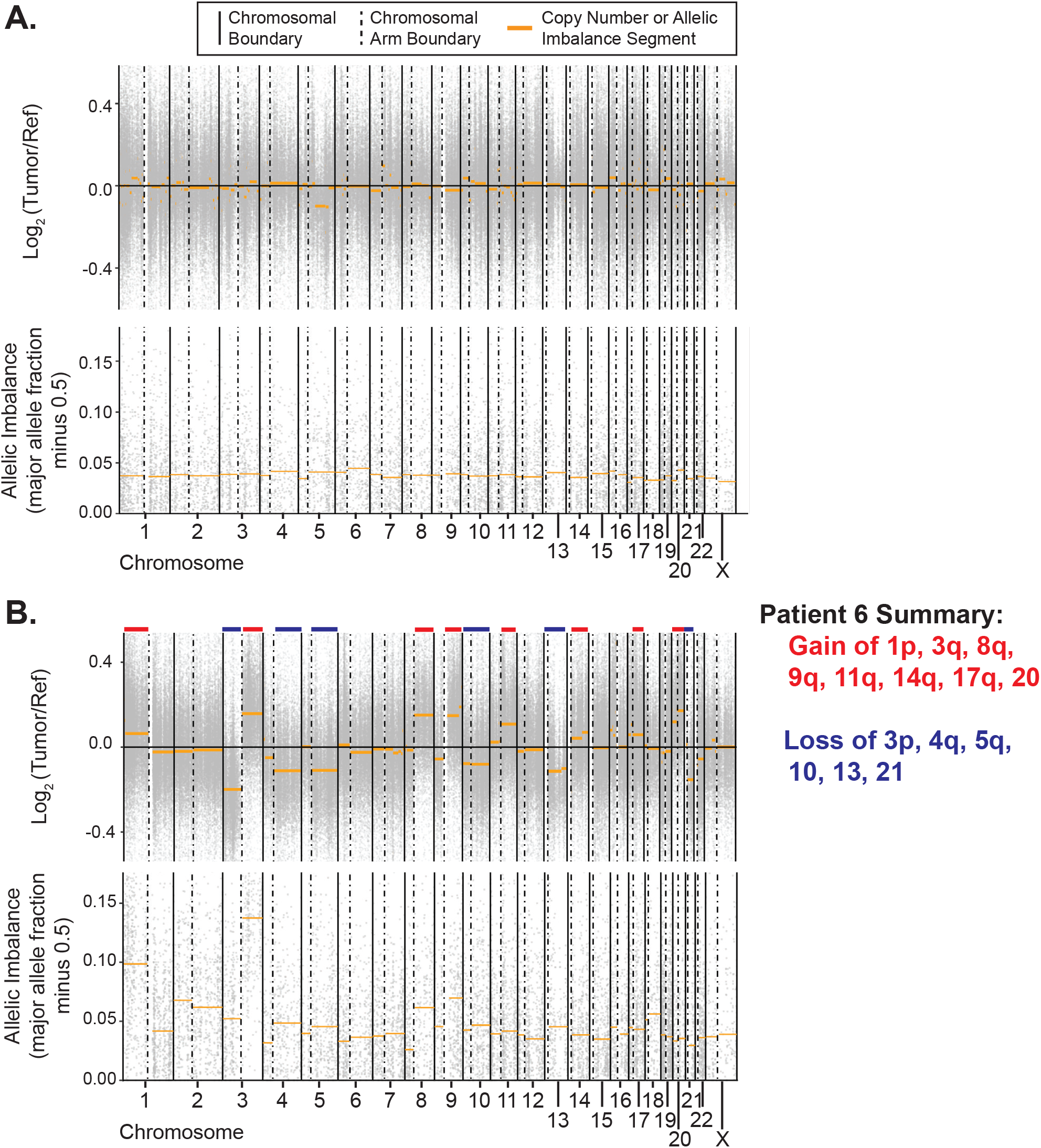
Copy number alterations and allelic imbalances in tumors from patients 4 and 6. **A-B.** Exome sequencing of DNA from bulk tumor tissue was performed from patients 4 (panel A) and 6 (panel B). *Top panel:* Copy number alterations were inferred over individual bins across the genome (grey data points) and segmented (gold lines) as described. *Bottom panel:* Allelic imbalance was inferred over germline heterozygous SNPs (grey data points) and segmented (gold lines) as described. In this plot, allelic imbalance equates to the major allele fraction minus 0.5. For example, a 50% to 50% or 60% to 40% ratio of reads, mapping to each allele of a heterozygous SNP, would respectively have imbalance values of 0.0 or 0.1 (i.e. the deviation from the expected fraction of 0.5). Overall, there were no compelling signals of copy number alterations or loss of heterozygosity in the patient 4 tumor, whereas several copy number alterations (noted) were present in the patient 6 tumor.

**Figure S2.**
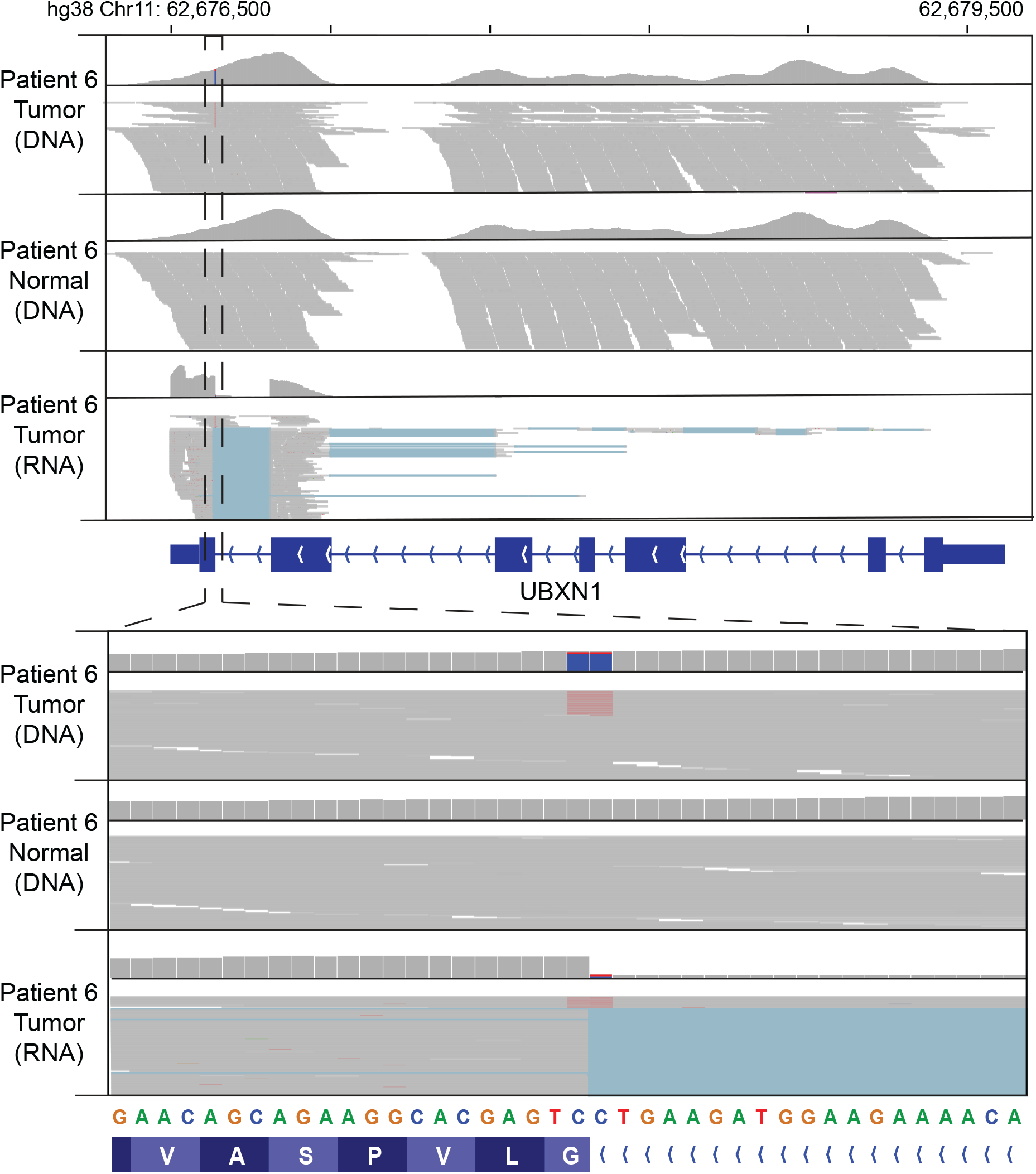
A splicing-site mutation affecting *UBXN1* is detectable in DNA- and RNA-sequencing data. Sequencing reads from exome sequencing of tumor DNA, exome sequencing of reference DNA, and RNA-sequencing of spatially barcoded cDNAs are visualized using the Integrative Genomics Viewer (IGV) browser. Reads are shown at gene-and exon-level of resolution, as indicated. Within each dataset, the upper track shows relative sequencing coverage and the lower track shows individual sequencing reads. Variant reads exceeding 10% allele frequency is colored. Note the bias in sequencing coverage toward the 3’ end of the gene in the spatial transcriptomic data. Also note how the mutant allele fails to properly splice.

**Figure S3.**
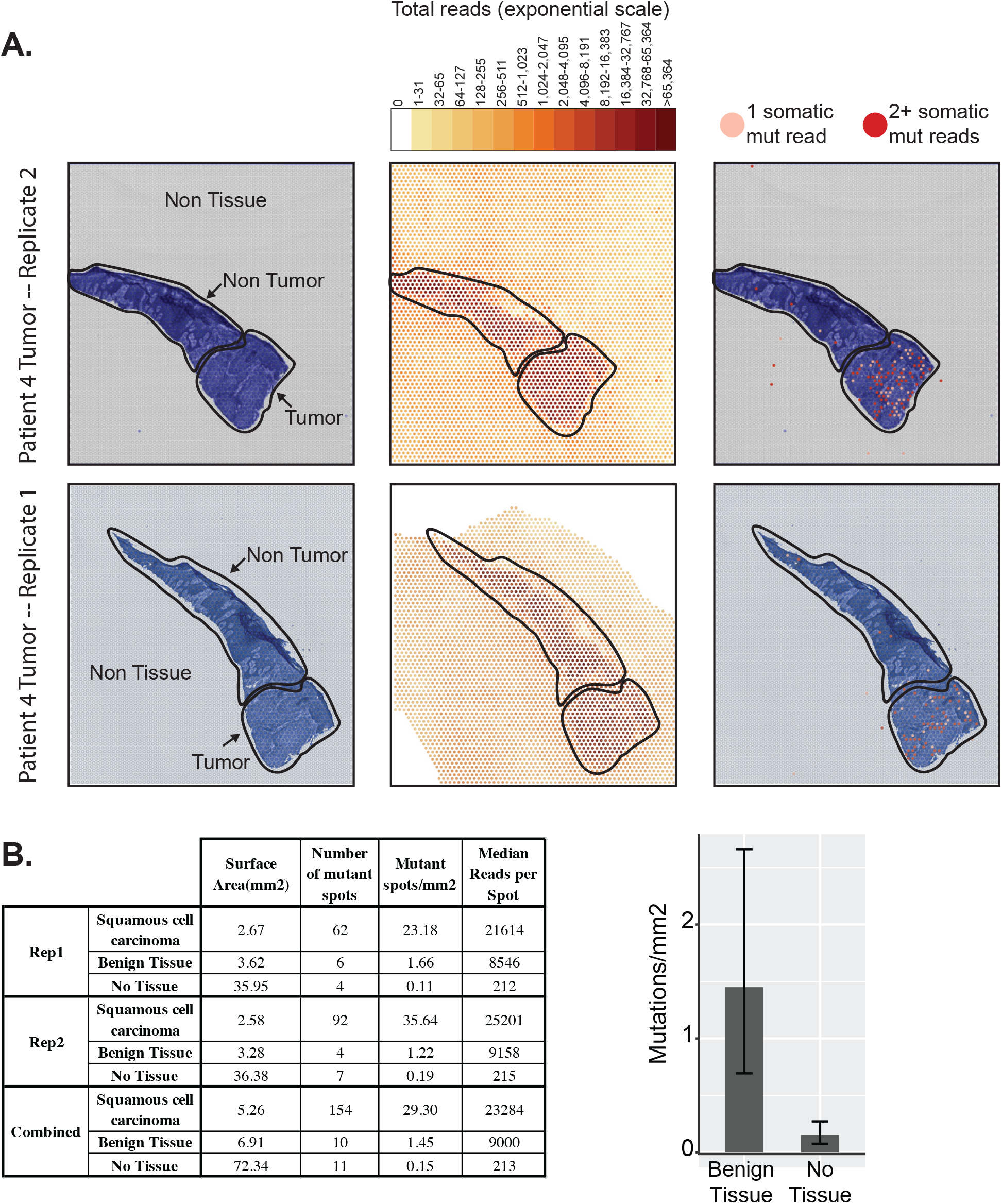
An excess of mutant reads in histologically benign tissue. Background signals were inferred by measuring total- and mutant-read counts in areas with no tissue, and these background signals were compared to the observed signals in spots overlaying tissue. **A.** The left panel marks the tumor, non-tumor, and non-tissue areas in the capture areas of each replicate from the patient 6 biopsy. The middle panel shows the number of reads per spot (note the exponential scale), and the right panel shows the spots with somatic mutation reads. **B.** The table on the left summarizes the surface area, number of mutant spots, mutant spot density, and reads per spot in each region of the two replicates as well as the combined data from the two replicates. The bar graph specifically highlights the mutation density in the benign tissue versus background (non-tissue spots) from the combined data. Error bars represent 95% confidence intervals using the Poisson test.

**Figure S4.**
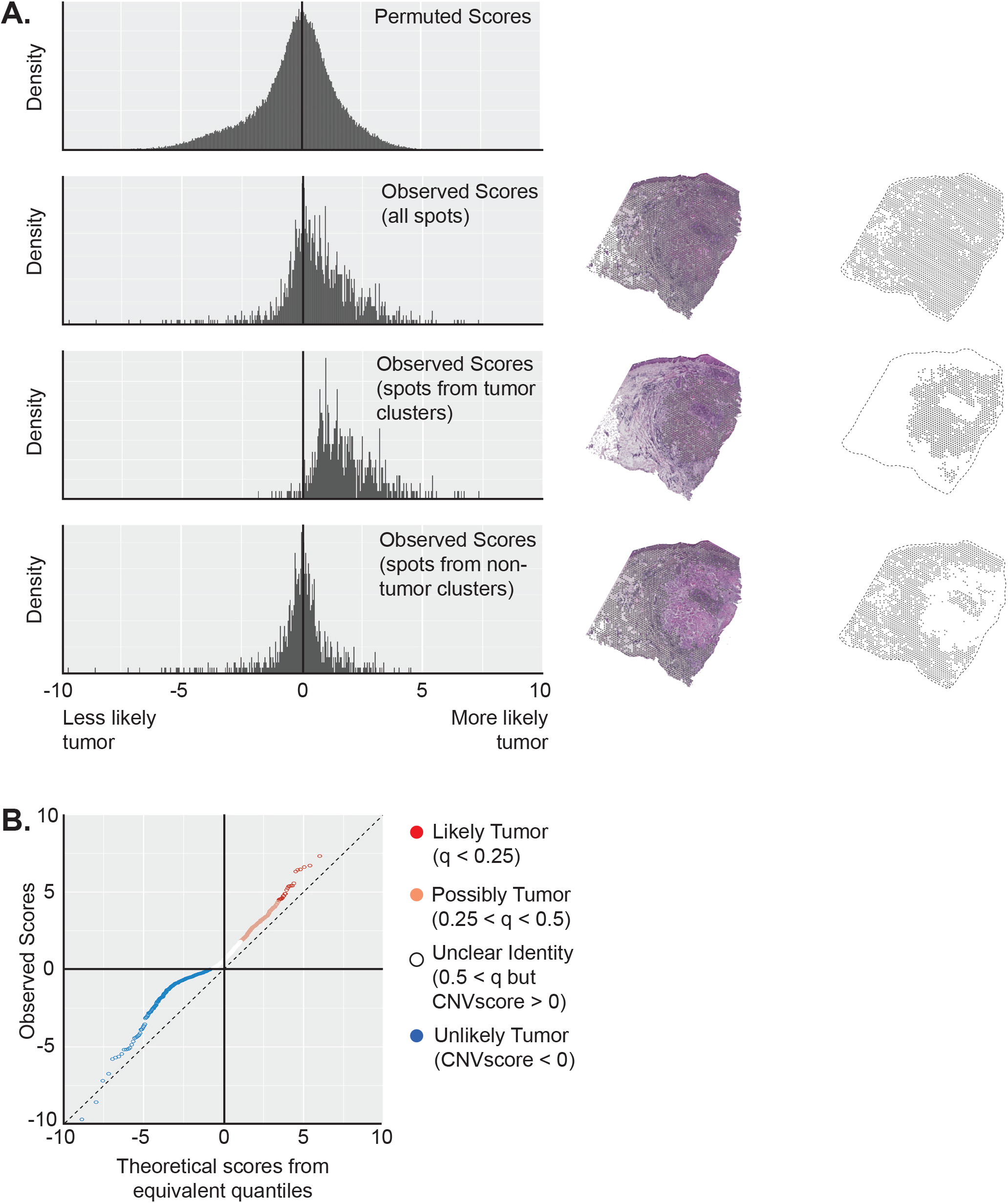
An enrichment of spots with copy number alterations. Copy number alterations were inferred from each spot’s RNA-sequencing data, as described. A “score” was calculated (see methods) to reflect how similar the RNA-inferred copy number profile of each spot was to the DNA-inferred copy number profile of the bulk tumor shown in Fig. S1 (spots with a higher score had copy number profiles that mirrored the DNA-inferred profiles). Scores were also calculated on permuted data to indicate the spectrum of scores that could occur by random chance. In panel **A**, histograms of scores are shown for permuted data, for all spots, and for subsets of spots, as indicated. Note that the permuted scores are centered at 0 while the observed histogram is shifted to the right. The rightward shift is driven by spots from gene expression clusters that are thought to derive from tumor cells, as shown in the subsetted data. In panel **B**, a quantile-quantile (Q-Q) plot compares the observed scores to equivalent quantiles from the permuted data. Note the off diagonal shift, confirming the skew in observed data towards higher scores. False discovery rates were calculated by comparing the frequency of permuted scores (false positives) to observed scores (total positives) and used to threshold the spots into 4 categories -- “Likely Tumor”, “Possibly Tumor”, “Unclear Identity”, or “Unlikely Tumor”.

**Figure S5.**
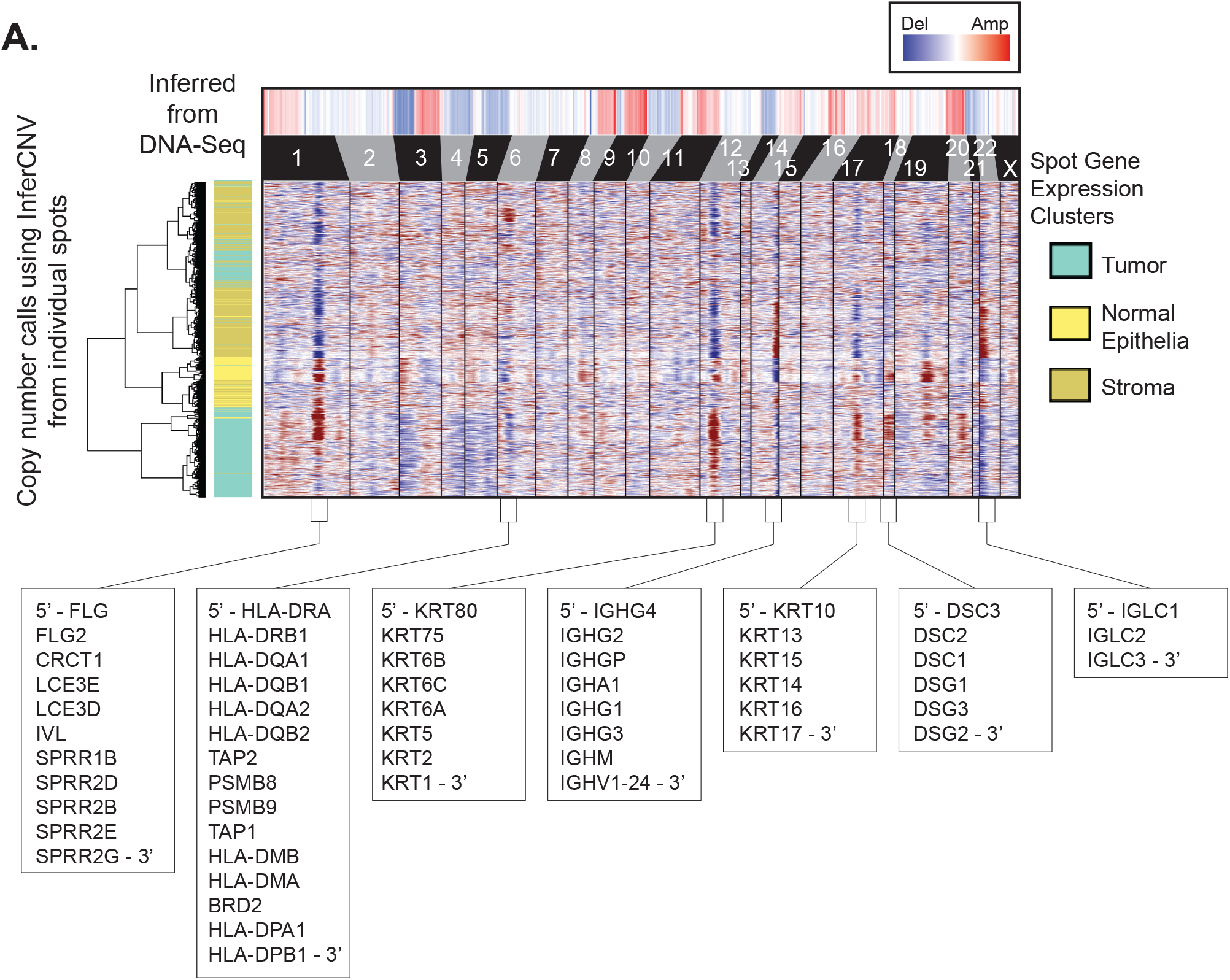
Neighborhoods of lineage-defining genes produce false positive calls when inferring copy number via a moving average of gene expression. Copy number alterations (CNAs) were inferred from DNA-Sequencing data (top heatmap) and from RNA-sequencing data of individual spots using InferCNV (lower heatmap). Spots (rows in the lower heatmap) are clustered based on the similarity of their copy number profiles. Spots were classified into histological categories based on their gene expression clusters and labeled as shown. Several of the most prominent copy number calls are highlighted with congtiguous genes within each region listed. Note how the highlighted copy number alterations center around lineage defining genes, whose relatively high expression in certain cell types likely produced false positive copy number inferences.

**Figure S6.**
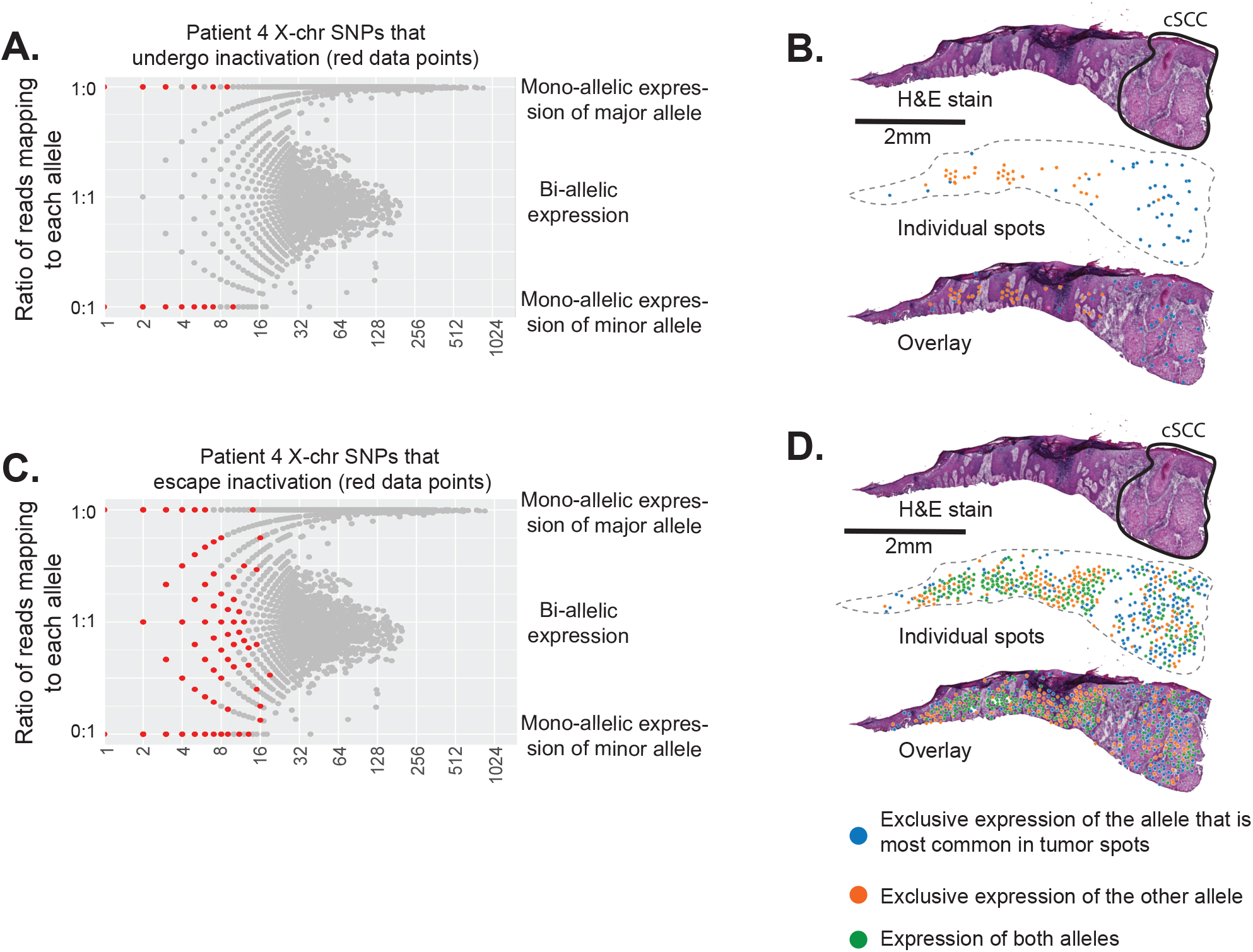
X-chromosome inactivation is detectable in spatial transcriptomics data. **A.** For each heterozygous SNP from each spot, the ratio of reads mapping to each allele is plotted as a function of total read coverage. Mono-allelic expression of X-chromosome SNPs was observed, presumably due to X-chromosome inactivation. Datapoints from SNPs known to escape X-chromosome inactivation were not highlighted in this plot and instead are shown in panel C. **B.** SNPs from panel A that were expressed in at least two tumor spots are projected onto spatial transcriptomic maps. These SNPs illustrate how different tumor spots tended to express the same allele (colored blue). Orange spots exclusively express the allele that was less often observed (typically never observed) in tumor spots. Spots expressing both alleles would be colored green, but no such spots exist for the SNPs that are subject to X-chromosome inactivation. Note how allelic expression of X-chromosome inactivated genes can illuminate mosaicism in tissues. **C-D.** Data for SNPs chrX:72273841 (C/T) and chrX:2782116 (G/A) (hg38 genome build), plotted as shown in panels A and B. These SNPs respectively reside in the *RPS4X* and *XG* genes, which are known to escape X-chromosome inactivation. Note the bi-allelic expression pattern in panel C and the admixing of spots expressing each allele (or both alleles) in panel D.

